# Degradation of mixed-linkage (1,3;1,4)-β-D-glucan in maize is mediated by the CAL1 licheninase

**DOI:** 10.1101/2020.11.16.385146

**Authors:** Florian J Kraemer, China Lunde, Moritz Koch, Benjamin M Kuhn, Clemens Ruehl, Patrick J Brown, Philipp Hoffmann, Vera Göhre, Sarah Hake, Markus Pauly, Vicente Ramírez

**Affiliations:** Department of Plant & Microbial Biology, Energy Biosciences Institute, University of California Berkeley, California, USA; Plant Gene Expression Center, Agricultural Research Service, U.S. Department of Agriculture, Albany, California, USA; Department of Crop Sciences, University of Illinois, Urbana, Illinois, USA; Institute of Microbiology/Group Pathogenicity, Heinrich Heine University Düsseldorf, Düsseldorf, Germany; Institute for Plant Cell Biology and Biotechnology – Cluster of Excellence on Plant Sciences, Heinrich Heine University Düsseldorf, Düsseldorf, Germany

## Abstract

The presence of mixed-linkage (1,3;1,4)-β-D-glucan (MLG) in plant cell walls is a key feature of grass species such as cereals - the main source of calorie intake for humans and cattle. Accumulation of this polysaccharide involves the coordinated regulation of biosynthetic and metabolic machineries. While several components of the MLG biosynthesis machinery have been identified in diverse plant species, degradation of MLG is poorly understood. A large-scale forward genetic maize screen for mutants with altered cell wall polysaccharide structural properties resulted in the identification of *candy-leaf1* (*cal1*). Cell walls of CAL1-deficient plants contain higher amounts of MLG in several tissues, including adult leaves and senesced organs, where only trace amounts of MLG are usually detected. In addition, *cal1* plants exhibit increased saccharification yields upon enzymatic digestion. Stacking *cal1* with lignin-deficient mutations results in synergistic saccharification increases. Identification of the causative mutation revealed that *CAL1* encodes a GH17 licheninase. Maize plants overexpressing CAL1 exhibit a 90% reduction in MLG content, indicating that CAL1 is not only required, but its expression sufficient to degrade MLG. CAL1 specifically hydrolyzes (1,3;1,4)-β-D-Glucans *in vitro*, and the single CAL1^E262K^ amino acid substitution is able to block all detectable activity. Time profiling experiments indicate that wall MLG content is modulated during day/night cycles inversely correlating with *CAL1* transcript accumulation. This cycling is absent in the *cal1* mutant, suggesting that the mechanism involved requires MLG degradation that may in turn regulate *CAL1* gene expression.

**One sentence summary** Mixed-linkage glucan is degraded by the CAL1 licheninase in maize

## INTRODUCTION

Plant cells are surrounded by a cell wall made of complex networks of various polysaccharides, phenolic compounds and proteins. A generic wall is based on cellulose microfibrils embedded in a matrix of various hemicelluloses, pectins, and/or lignin. However, the specific components, their relative abundance and their biochemical interactions are specific to the plant species and the cell types (Pauly and Keegstra 2010). Plant cell walls have been classified as type I wall (dicots and non-commelinid monocots) and a structurally different type II wall found in grasses (Poaceae) (Carpita and Gibeaut 1993; Somerville et al. 2004). In type II walls, the main hemicellulose found cross-linking cellulose microfibrils is glucuronoarabinoxylan (GAX), composed of a substituted β1,4-linked xylose backbone. Furthermore, compared to type I walls, grass walls contain very low amounts of xyloglucan, pectic polysaccharides and structural proteins. Also, type II wall lignin contains ρ-hydroxyphenyl units that are only found in trace amounts in type I walls (Grabber et al., 2004). Unlike a type I wall, grass walls also contain high levels of hydroxycinnamates such as ferulic acid or *p*-coumaric acid, covalently linked to GAX and lignin (Saulnier et al., 1999, Hatfield et al., 1999). Another major difference of type II walls is the occurrence of mixed-linkage glucan (MLG), an unsubstituted homopolymer where β-1,4-linked glucose oligomers are connected by β-1,3 linkages (Carpita and Gibeaut, 1993; Carpita, 1996). The length and relative proportions of the oligomers varies among species but usually cellotrioses and cellotetraoses are dominant (Staudte et al., 1983). While MLG has not been observed in type I walls, it has been found in the walls of unrelated organisms such as ancient vascular plants, green algae, liverworts, bacteria, fungi, and lichens (Gorin et al., 1988; Stone and Clarke, 1992; Popper and Fry, 2003; Fry et al., 2008; Pettolino et al., 2009; Honegger and Haisch, 2001), suggesting independent evolution of this polysaccharide in multiple species (Fincher, 2009).

Advances have been made regarding the mechanism of MLG biosynthesis through the discovery and characterization of several cellulose synthase-like (CSL) proteins. Heterologous expression of various plant CSLF, CSLH and CSLJ proteins result in the production and deposition of MLG in the walls of non-grass species that usually lack MLG (Burton et al., 2006; Doblin et al., 2009; Little et al., 2018). Multiple lines of evidence indicate that CSLF6 plays a major role in MLG biosynthesis in most grass species. CSLF6 transcripts are the most abundant among the MLG biosynthetic genes in barley, wheat and Brachypodium. Also, CSLF6 overexpression leads to increased MLG levels in transgenic plants, while CSLF6 loss-of-function results in a strong reduction in MLG abundance (Burton et al., 2008; Christensen et al., 2010).

MLG is considered a plant stage-specific polysaccharide and its accumulation appears to be regulated by synchronized mechanisms of biosynthesis and degradation. Several grass species accumulate large amounts of MLG (up to 20% dry mass) in actively growing tissues, and then degrade it once the growth/cell elongation stage ends (Carpita 1996; Kim et al 2000; Carpita and McCann 2010). Due to its transitory nature, MLG has been considered a storage molecule for rapid carbon release in the growth phase of cells (Roulin et al., 2002). A handful of degrading enzymes with (1,3) (1,4)-β-D-glucan-4-glucanohydrolase activity or licheninases, have been identified in barley, wheat, and rice and their respective biochemical activities characterized (Litts et al., 1990; Wolf et al., 1991; Lai et al., 1993). However, their specific biological role is largely speculative.

In the last 20 years, forward genetics has been used to identify mutants with altered plant cell walls. A myriad of large-scale screens have been performed, reaching near-saturation in model species such as *Arabidopsis thaliana*, a type I wall plant (For example: Reiter et al., 1997; Turner and Somerville, 1997; Chen et al., 1998; Gille et al., 2009). These screens have been extremely useful in deciphering the mechanisms involved in the biosynthesis, deposition, turnover, and function of wall components. However, the number of high-throughput mutant screens performed in grass species is significantly lower (Carpita and McCann, 2002; Marriott et al., 2014). As a result, the catalog of wall-altered grass mutants is largely incomplete, and grass-specific aspects of the wall, such as MLG biology, remain underexplored.

Here, a forward genetic screen was performed on a chemically mutagenized maize population, designed to identify mutants with altered cell wall structures and/or properties. We identified and characterized *cal1*, to our knowledge the first loss-of-function mutant impaired in MLG degradation in a grass, here maize. In addition, we provide insights into a poorly understood mechanism regulating MLG turnover following a circadian rhythm.

## RESULTS

### Identification and characterization of the maize *candy-leaf1* (*cal1*) mutant

An EMS-mutagenized maize population was screened for individuals with altered cell wall attributes. For this purpose, destarched alcohol-insoluble residue was prepared from second leaves of individual 2-week-old mutagenized maize plants and its monosaccharide composition was determined. One of the mutants, termed *candy-leaf1* (*cal1*), was identified because compared to the A619 non-mutagenized wild-type control, its wall material exhibited a 248.5 ± 8.5 % increase in wall glucose (Glc) upon acid hydrolysis. In contrast, the content of two other abundant monosaccharides measured -xylose (Xyl) and arabinose (Ara)- remained unaltered (**Fig. 1A**). In addition, the high glucan content was accompanied by a 30.7 ± 6.2 % increase in enzymatic saccharification yield compared to the wild type (**Fig. 1B**). A more detailed wall analysis of *cal1* indicated no difference in cellulose abundance, wall-bound *O*-acetate, lignin content or relative monolignol composition (**Fig. 1C**). The increased *cal1*-associated glucan content initially observed in young seedlings was also found in adult leaves (leaf above the ear), senesced flag leaves and senesced stalks (internode above the ear), with increases of 51.3 ± 9.7, 62.3 ± 11.2 and 27.9 ± 7.6% in Glc content compared to A619 wild type, respectively. Similarly, saccharification yields were also significantly increased in adult and senesced leaves from *cal1* individuals (**Table S1**).

**Figure 1.**
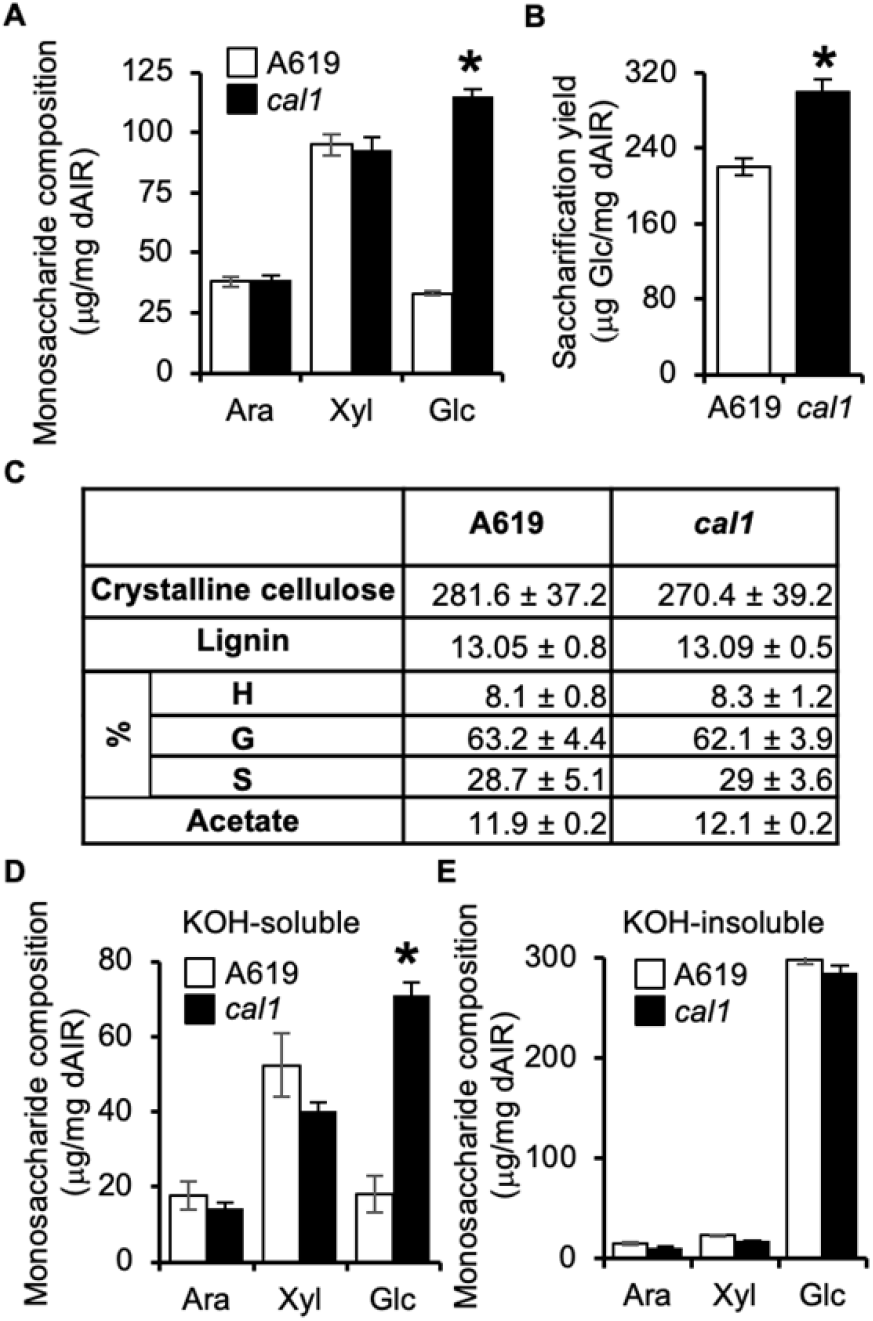
Cell wall composition of the maize cal1 mutant. (A) HPAEC-based quantification of monosaccharides released by acid hydrolysis. (B) Glc released by enzymatic digestion of dAIR. (C) Table summarizing the content in A619 and cal1 in terms of crystalline cellulose (μg/mg dAIR), lignin content (% of acetyl bromide soluble lignin) and lignin composition (%), and wall-bound 0-acetate (μg/mg dAIR). (%) means lignin composition as the distribution of guaiacyl (G), p-hydroxyl phenol (H) and syringyl (S) monolignols. No significant differences were found between genotypes (Student’s t-test, p<0.05). (D-E) HPAEC-based quantification of the monosaccharides present in KOH-soluble (D) and KOH-insoluble (E) fractions. All bar diagrams represent the mean± SD of 4 biological replicates. A619 (white bars) and cal1 (black bars). Asterisks indicate statistical difference between the two genotypes according to Student’s t-test (p<0.05).

To further investigate the details of *cal1*-associated wall alterations, an alkali extracted hemicellulosic fraction of *cal1* and wild-type A619 walls was subjected to quantitative monosaccharide composition analysis. This KOH-soluble, hemicellulose-enriched *cal1* fraction showed a 293.8 ± 17.1 % increase in Glc content compared to A619 (**Fig. 1D**). In contrast, no differences were found in the abundance of Glc, Xyl or Ara in the remaining KOH-insoluble residue (**Fig. 1E**). These results suggest that a Glc-containing hemicellulose polymer is likely affected in *cal1* rather than cellulose. In order to determine its identity, glycosyl-linkage analysis was performed on the KOH-soluble fraction. An increase in the relative abundance of 3- and 4-linked glucopyranosyl (Glcp) residues in the *cal1* hemicellulose fraction was observed when compared to A619 indicative of a mixed-linkage glucan (MLG), while the remaining linkages were slightly decreased or unchanged (**Fig. 2A**).

**Figure 2.**
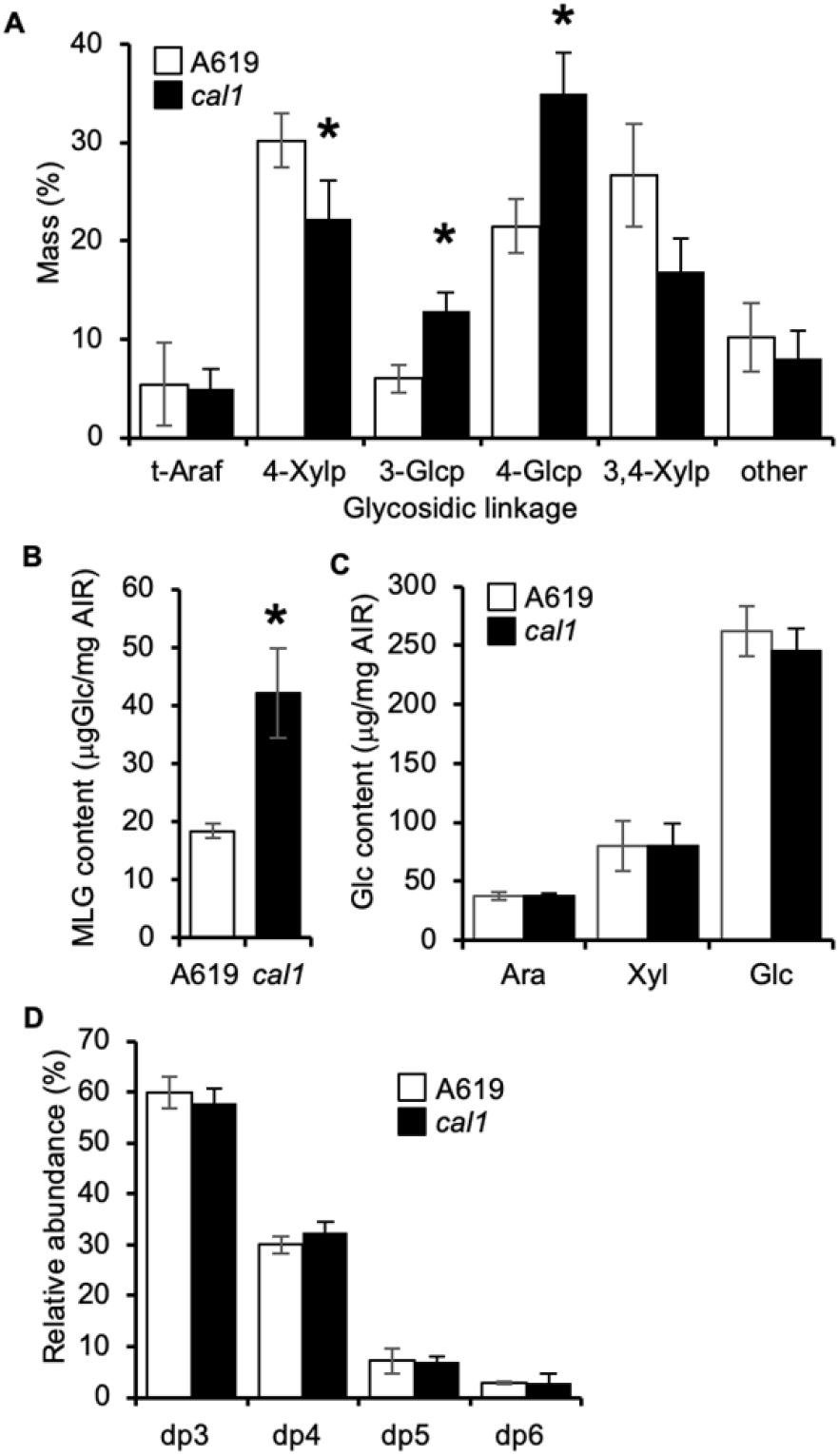
MLG content and structure in *cal1*. (A) Glycosidic linkage analysis of released oligosaccharides after licheninase treatment of AIR. Values represent the percentage of the total ion chromatogram peak area of the indicated partially methylated, acetylated alditols. t-Araf= terminal arabinofuranosyl; 4-Xylp= 4-linked xylopyranosyl; 3-Glcp= 3-linked glucopyranosyl; 4-Glcp= 4-linked glucopyranosyl; 3,4-Xylp= 3,4-linked xylopyranosyl. (B) MLG content measured as Glc released after licheninase digestion of AIR, followed by glucosidase treatment of the resulting β-glucan oligosaccharides. (C) Glc content released by Saeman hydrolysis of the insoluble residue left after licheninase treatment in (B). (D) HPAEC-based quantification of β-glucan oligosaccharide products released by licheninase treatment of AIR. dp=degree of polymerization of detected hexose oligomers. All bar diagrams represent the mean ± SD of 4 biological replicates. A619 (white bars) and cal1 (black bars). Asterisks indicate statistical difference between the two genotypes according to Student’s I-test (p<0.05).

Hence, *cal1* walls seem to have a higher abundance of MLG. To explore this hypothesis, the amount of MLG in the plant tissue was determined using a specific (1,3) (1,4)-β-D-glucan endoglucanase (licheninase). Digestion of AIR with this enzyme results in the release of β-glucooligosaccharides specifically derived from MLG that can then be hydrolyzed into Glc following glucosidase treatment (McCleary and Glennie-Holmes, 1984). Using this method, *cal1* walls showed a 229.2 ± 41.9 % increase in MLG content compared to A619 (**Fig. 2B**). Further analysis of the licheninase-resistant residue showed no differences in monosaccharide composition among the genotypes (**Fig. 2C**). Combined, these results confirm that the high glucan content associated with *cal1* results from a hyperaccumulation of MLG.

The β-glucooligosaccharides generated by licheninase treatment of AIR from *cal1* and A619 were also quantified. Mainly hexose oligomers with degrees of polymerization (DP) of 3 and 4 were present, although small amounts of oligomers with a DP5 and DP6 were also detected (**Fig. 2D**). No differences between *cal1* and A619 were found in the relative distribution of the β-glucooligosaccharides indicating that MLG accumulates in *cal1* walls, but its structure is unaltered.

### Effect of altered lignin on the increased saccharification yield associated with *cal1*

One hypothesis is that the enhanced saccharification yield observed in *cal1* is caused by a high MLG content. No additional wall compositional differences were detected in *cal1* walls, including crystalline cellulose content or the amount and composition of lignin, both factors reported to strongly influence saccharification yield (Grabber 2005; Himmel et al., 2007). Lignin negatively impacts saccharification efficiency by hindering the action of hydrolytic glycanases and glycosidases. Accordingly, reduced lignin content and changes in the lignin structure, as those reported in some *brown-midrib* (*bm*) mutants, have a positive influence on wall digestibility (Saballos et al., 2008; Dien et al., 2009; Xiong et al., 2019). To study the effect of altered lignin content and composition in the enhanced saccharification yield exhibited by *cal1*, the *cal1* mutant was crossed with individual *bm* lines affected in the lignin biosynthetic pathway, and the resulting double mutants were analyzed. Maize *bm1* and *bm3* encode a cinnamyl alcohol dehydrogenase (CAD) and caffeoyl-O-methyltransferase (COMT), respectively (Halpin et al., 1998, TPJ; Vignols et al., 1995). Comparative analyses of the saccharification yield and related wall composition traits (i.e. glucan content, total lignin content and lignin composition) were performed (**Fig. 3** and **Table S3**). As previously observed, *cal1* exhibited increased saccharification values (≈25%). Interestingly, under the experimental saccharification conditions used here, no differences were observed for the saccharification yield of individual *bm1* and *bm3* single mutant lines compared to the B73 control. However, in the case of the *cal1 bm1* and *cal1 bm3* double mutants, 51% and 59% saccharification yield increases were noted compared to the wild type, respectively, suggesting a synergistic effect on saccharification efficiency between *cal1* and each of the *bm* mutations used (**Fig. 3B**). The two *cal1 bm* double mutants and *cal1* single mutant exhibited similar increases in wall Glc content compared to the wild type confirming the high MLG abundance (**Fig. 3A**). In terms of lignin content only a slight reduction in lignin content was observed in the *cal1 bm1* line but not in the single mutant lines likely due to the developmental stage of the plants analyzed, i.e. second leaf of only 2-week-old seedlings. However, both *bm3* and *cal1 bm3* plants showed increased relative amounts of guaiacyl (G) and decreased syringyl (S) monolignol units (**Table S3**). These results suggest that the increased saccharification yield observed in the *cal1 bm*1 and *cal1 bm3* lines compared to *cal1* are not correlated with differences in lignin or MLG contents. Instead, in the *cal1 bm* double mutant lines the combined increase of MLG content and altered lignin may lead to a different wall architecture resulting in the observed synergistic enhancement of wall digestibility.

**Figure 3.**
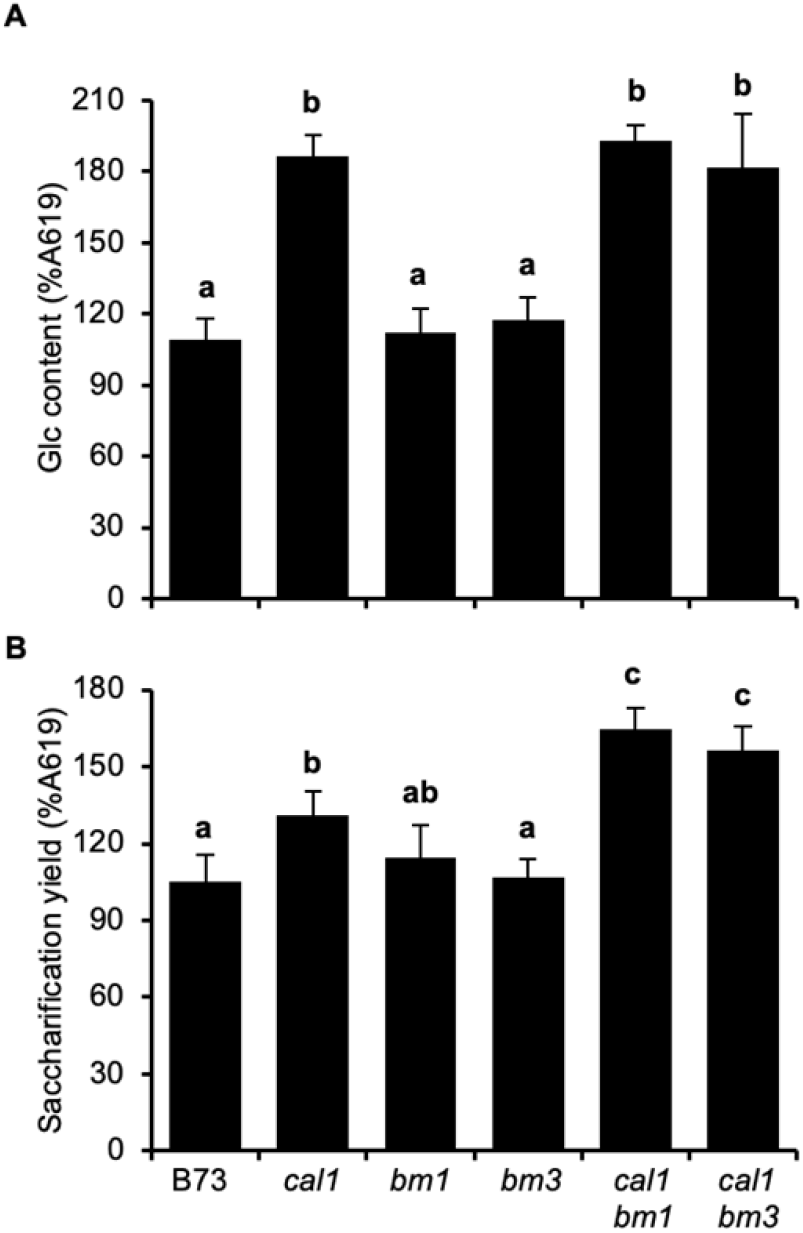
Effect of altered lignin on *cal1* phenotype. (A) Glc content of the hemicellulosic wall fraction (TFA hydrolysis). (B) Glc released by enzymatic digestion of dAIR. All bar diagrams represent the mean ± SD of 4 biological replicates. Different letters indicate statistical differences based on ANOVA analysis with post-hoc Tukey Honest Significance Difference test.

### Effect of *cal1* on plant growth and interaction with a pathogenic fungus

A detailed analysis of the seedling growth habit under controlled conditions did not reveal any difference between *cal1* and the non-mutagenized control (**Fig. S1**). When grown in the field, no significant differences were found in overall plant architecture between wildtype and *cal1* plants (**Fig. S2)**. Diverse plant fitness-related traits were also analyzed (**Table S2**). No differences were found in the moisture content in whole above-ground plant biomass or grain between the *cal1/cal1* mutant in an inbred A619 background and wild-type (*CAL1/cal1*) Mo17/A619 or B73/A619 hybrids. Only marginal differences in grain dry weight were found between the wild-type Mo17/A619 hybrid and the *cal1/cal1* mutant (p=0.04) and differences in biomass dry weight between these two were not significant (p=0.46). As expected, the B73/A619 hybrid, which is a much more heterotic combination than Mo17/A619 (Melchinger et al., 1991), yielded significantly more grain and biomass than either of the other two entries. Overall, these results indicate no major impact of the *cal1* mutation on plant development and reproduction.

A higher abundance of a more easily degradable glucan in the walls of *cal1* might lead to a higher susceptibility of plant pathogens. This does not seem to be the case. When *cal1* and wild-type seedlings were challenged with the pathogenic fungus *Ustilago maydis,* no differences were found on symptom development in terms of tumor formation, anthocyanin accumulation or appearance of chlorotic lesions (**Fig. S3**).

### Mapping of *cal1* mutation

To identify the causative mutation responsible for the *cal1* chemotype, a mapping population was obtained by crossing *cal1* in the A619 inbred by the Mo17 inbred and self-pollinating that hybrid. The glucan content of 101 individuals of the F2 mapping population was determined. From those, 25.25% of the individuals displayed a ≥200% increase in glucan content compared to A619, consistent with a recessive mutation. A Chi-square statistical analysis supported *cal1* being a recessive mutation (**Fig. 4A**). The causative mutation was mapped to an interval of 10.46Mb on Chromosome 6 (**Fig. S4)**. After sequencing of selected candidates among the gene models in this region (**Table S4**), a G to A change was found in the second exon of gene *GRMZM2G137535* causing a putative glutamic acid (E) to lysine (K) conversion in the encoded protein (**Fig. 4B**). A BLAST search resulted in the identification of multiple proteins with high sequence similarity and identity to GRMZM2G137535 within grass species. Phylogeny analysis showed that GRMZM2G137535 clusters together with proteins with predicted licheninase activity from the Glycoside Hydrolase Family 17 (GH17) according to CAZy classification (www.cazy.org; **Fig. S5**). Among them, GRMZM2G137535 shares 86.6, 84.9 and 86.3% identity with biochemically characterized licheninases isolated from germinated barley (EI and λHv29) and wheat (λLW2) grains, respectively (Litts et al., 1990; Wolf et al., 1991; Lai et al., 1993). The E262K substitution in *cal1* is located in a residue highly conserved among all four proteins (**Fig. 4C**). Moreover, the glutamic acid E262 is predicted to be a part of the catalytic triad required for catalytic activity of the enzyme. Besides the barley and wheat putative homologs, other GH17 proteins are found in the same cluster with high identity (>80%) to GRMZM2G137535 encoded by the sorghum, oat, rice, *Brachypodium distachyon*, *Panicum hallii* and *Setaria italica* genomes (**Fig. S5**). These results suggest that GRMZM2G137535 may encode a licheninase highly conserved among grasses.

**Figure 4.**
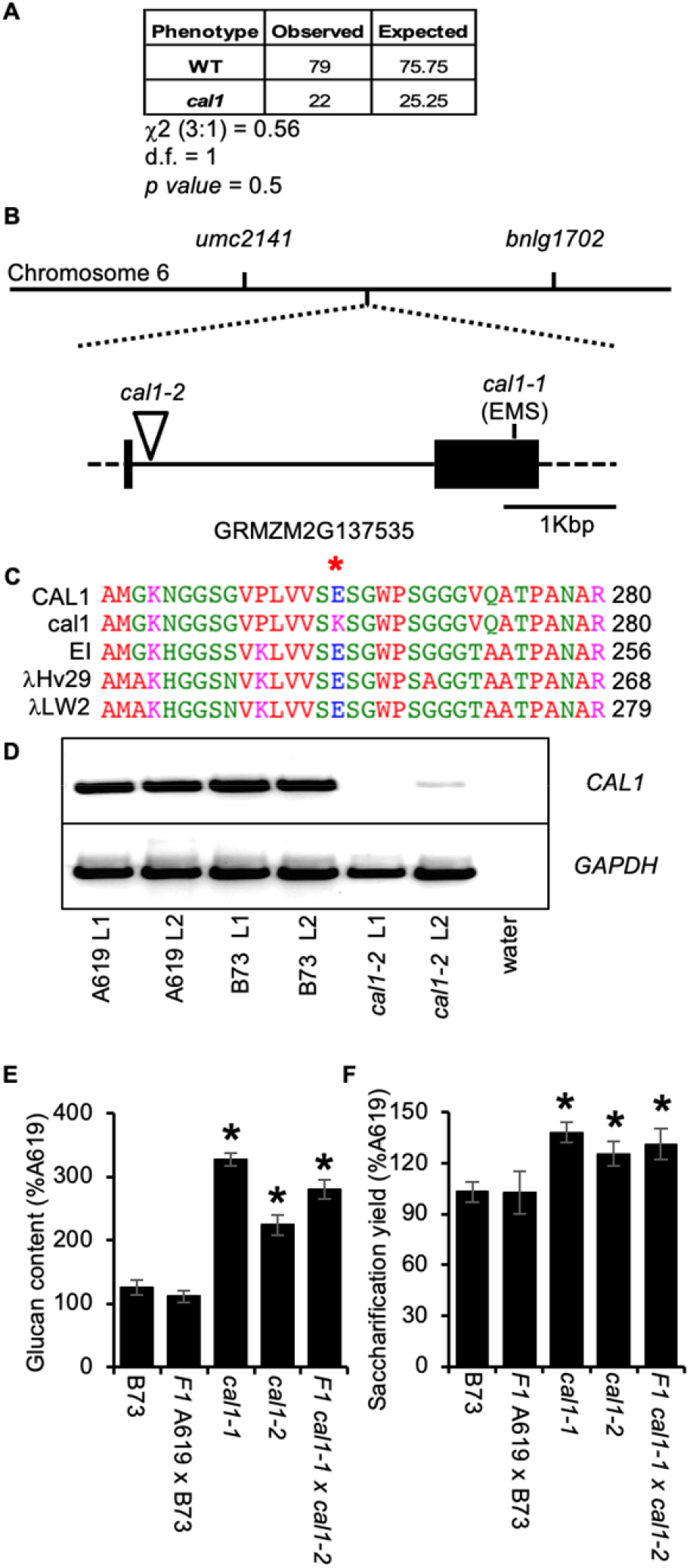
Two allelic mutants affected in the maize CAL 1 grass-specific gene encoding a putative licheninase. (A) Chi-square (χ^2^) test for a recessive segregation (3: 1) of *cal1-1* in a F2 cross to B73. d.f.= degree of freedom. p values>0.05 means hypothesis must be accepted. (B) Schematic representation of cal1-1 mapping on chromosome 6 between the *umc2141* and *bnlg1702* markers, and structure of the *CAL 1* gene. Black boxes= exons; lines= introns; dashed lines= untranslated regions. Position of the cal1-1 and cal1-2 mutations are indicated. (C) Detail of the protein alignment of maize CAL1, cal1-1 and three highly similar licheninases from barley (El and λHv29) and wheat (λLW2). Red asterisk indicates the mutated amino acid residue in *cal1-1*, an E262K substitution. Numbers to the right indicate the position of the last aminoacid residue shown (R) related to the starting methionine. (D) Transcript level of *CAL1*. RT-PCR with CAL 1-specific primers using cDNA synthetized from RNA extracted from the first (L 1) and second (L2) leaf of the indicated genotypes. A negative control was included using water as template. Amplification with GAPDH-specific primers was used as housekeeping control. (E) Quantification of the hemicellulosic Glc released after TFA hydrolysis of dAIR material from the indicated genotypes. (F) Glucose released by enzymatic digestion of dAIR from the indicated genotypes. Bars in E and F show the percentage of the A619 value. Values represent mean ± SD of 5 biological replicates. Black asterisks indicate statistical differences to B73 according to Student’s t-test (p<0.05).

A second *Mutator* allele, *cal1-2*, was found in the Trait Utility System for Corn (TUSC) collection developed by Pioneer Hi-Bred International (Meeley and Briggs 1995). The *cal1-2 Mutator* line contained an insertion in the +178bp position into the first intron, causing a reduction in GRMZM2G137535 transcript levels in the first and second leaves (**Fig. 4D**). Both glucan content and saccharification yield were increased in *cal1-2* compared to the wild-type control, although to a lesser extent than in the EMS *cal1* allele (now termed *cal1-1*). F1 plants generated by crossing *cal1-1* to *cal1-2* exhibited a mutant phenotype (i.e. increased glucan content and saccharification yield), indicating that the mutants are allelic (**Fig. 4E and 4F**). Together these results indicate that *CAL1* is *GRMZM2G137535*.

### CAL1 overexpression in maize plants

CAL1 overexpression (CAL1ox) lines were generated with increased *CAL1* transcript levels (**Fig. S6A**). Three independent CAL1ox lines were selected and crossed to B73 and the resulting F1 self-pollinated. In the segregating F2 populations, glucan content and saccharification were determined. Homozygous *cal1-1* plants were used as a control, exhibiting a 256% increase in glucan amounts compared to A619. After individual genotyping, plants negative for the presence of the CAL1ox transgene displayed a glucan content ranging from 98 to 123.8% of the A619 value, while CAL1ox-containing plants showed significant reductions, showing values from 49.0 to 59.4% compared to A619 (**Fig. S6B**). Similarly, saccharification yields were also reduced by 27.6 to 42.7% correlating with the presence of the CAL1ox transgene, in contrast to the CAL1ox-negative plants exhibiting values similar to A619 (106 to 116.4%). As expected, the *cal1-1* control showed an increased saccharification yield (130.5%) (**Fig. S6C**). These results indicate that overexpression of *CAL1* results in low MLG content and decreased enzymatic saccharification yield, the opposite chemotype in the *CAL1* loss of function alleles *cal1-1* and *cal1-2*.

### Characterization of CAL1 enzymatic activity

Wild-type (CAL1) and mutant (CAL1^E262K^) C-terminal His-tagged recombinant protein versions were produced in *Nicotiana benthamiana* by an *Agrobacterium tumefaciens*-mediated transient expression system. After affinity purification the licheninase activity of both recombinant proteins was assayed using commercial MLG from barley flour as substrate (**Fig. 5B**). Laminarin and cello-oligosaccharides were also used as potential substrates to test the β-1,3- and β-1,4-glucanase activities, respectively. Under the conditions used, CAL1 was only able to degrade MLG indicating its specific licheninase activity. The activity was abolished in the CAL1^E262K^ mutant version, indicating that the E262 residue is indeed required for protein activity and suggesting that *cal1-1* is a loss-of-function allele (**Fig. 5B**).

**Figure 5.**
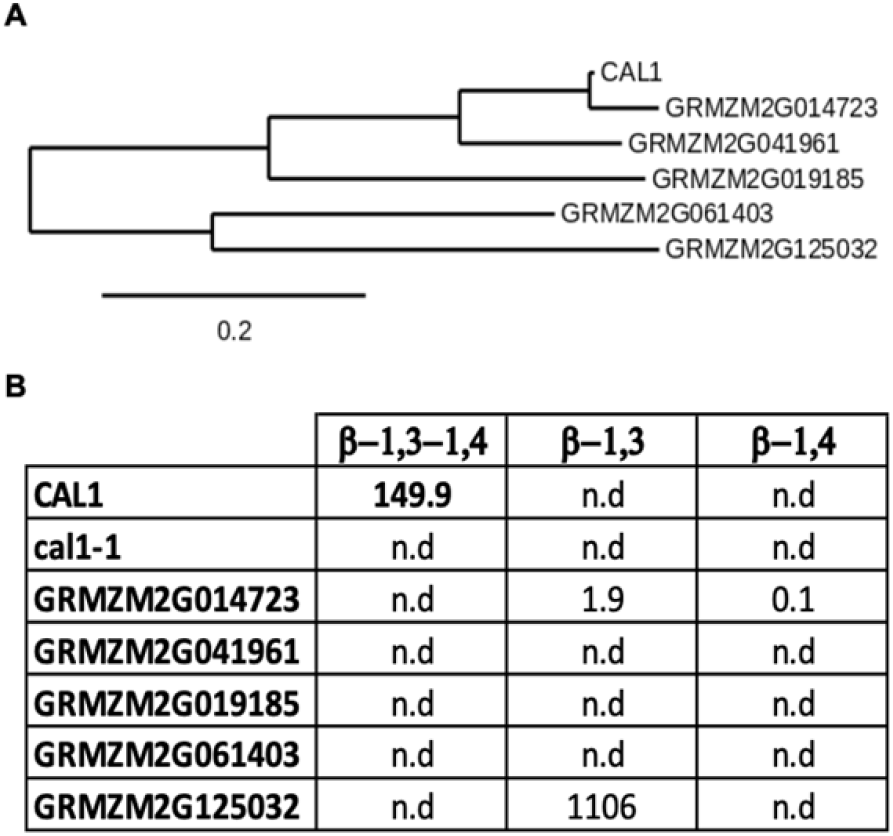
In vitro licheninase activity test of maize CAL 1 paralogs. (A) Detail of the maximum likelihood phylogenetic tree of CAL 1 and its closest para logs in maize. See Figure S7 for the complete tree. (B) Table showing the activity of CAL 1, CAL1^E262K^ mutant version (cal1-1) and 5 selected maize paralogs. β-1,3-1,4: licheninase activity using MLG as substrate. β-1,3: 1,3-β-glucanase activity using laminarin as substrate. β-1,4: 1,4-β-glucanase activity using cellooligosaccharides as substrate. Data shown in nKaVmg. n.d= no activity was detected

A survey of proteins encoded in the maize genome with sequence similarity to CAL1 resulted in 66 matches with identities ≥30%. From those, 23 proteins belong to GH17 and 43 to GH1 (**Fig. S7**). Only CAL1 and the three closest paralogs - two GH17 proteins (GRMZM2G014723 and GRMZM2G041961) and one GH1 protein (GRMZM2G019185) - have predicted licheninase activity according to the CAZy database, while the others were classified as glucan endo-1,3-β-glucosidases (34 total, 11 GH17 and 23 GH1), *O*-glycosyl hydrolases (18 total, 9 GH17 and 9 GH1) or with unknown function (10 total, 1 GH17 and 9 GH1). The 5 closest proteins to CAL1 were selected to test for potential licheninase activity, including the 3 mentioned putative licheninases (GRMZM2G014723, GRMZM2G041961 and GRMZM2G019185) and two other GH17 members with predicted glucan endo-1,3-β-glucosidase activity (GRMZM2G061403 and GRMZM2g125032; **Fig. 5A**). In contrast to CAL1, none of the recombinant proteins assayed were able to degrade MLG under the assay conditions tested and only GRMZM2g125032 showed 1,3-β-glucosidase activity (**Fig. 5B**). This *in vitro* data suggests no apparent genetic redundancy of the maize CAL1 licheninase explaining the observed drastic MLG chemotype in the *cal1-1* mutant.

### Dark-induced degradation of MLG

The accumulation of MLG in CAL1-deficient and CAL1-overexpressing plants compared to wild type was further investigated over a 48-hour period. Plants were grown in a 12h day, 24°C/12h, 20°C night photoperiod and the second leaf blade of 2-week-old plants were collected at different timepoints. As shown in **Fig. 6**, MLG content in wild-type plants progressively increases during the day hours, peaking at approximately 6h after the light cycle begins and remaining constant until dark. During the dark period, MLG content decreased, reaching minimum values after approximately 8h. In contrast, MLG content in *cal1-1* samples did not fluctuate but remained constant over the 2 day/night cycles studied here, with values similar to the maximum observed in wild-type samples. On the other hand, MLG content in CAL1ox plants remained low at all timepoints, with only 10% of the highest content found in wild-type plants at the end of the day.

**Figure 6.**
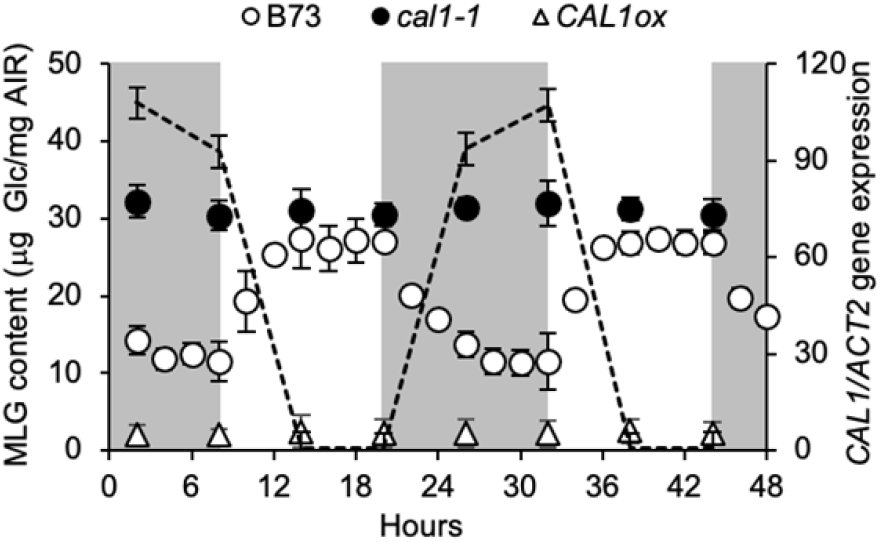
Time course of CAL 1 gene expression and MLG content. Left axis shows MLG content measured as Glc released after licheninase digestion of AIR followed by glucosidase treatment of the resulting β-glucan oligosaccharides. Wildtype (B73, empty circles), *cal1-1* mutant (black circles) and CAL 1ox (empty triangles) values shown as mean ± SD of 5 biological replicates. Dashed line shows the *CAL 1* gene expression in B73 measured by quantitative RT-PCR (right axis) as mean ± SD of 3 biological replicates. ACT2 gene expression was used for normalization. Samples for both analyses were taken at the indicated time points. Gray/white areas indicate subjective night and day respectively

The fluctuation in MLG content in wild-type plants was inversely correlated with *CAL1* gene expression (**Fig. 6**). *CAL1* expression was at a maximum around the end of the dark period, and its transcription was rapidly downregulated once the light cycle started. A 20000-fold night-triggered *CAL1* transcript upregulation was observed comparing the values at the end of the night period (8 and 32h) with the end of the day (20 and 44h). Thus, *CAL1* transcription was maximal when MLG levels are lowest in a circadian rhythm.

As an alternative energy store in plants starch turnover was also investigated in CAL1-deficient or CAL1-overexpression plants. Starch is the most abundant carbohydrate reserve in plants and is accumulated during the day and broken down at night (reviewed in Zeeman et al., 2010). Both starch and MLG contents were quantified in 2-week-old plants after 8 and 16 hours of being transferred to dark conditions and compared to plants kept in constant light. As shown in **Fig. S8**, degradation of MLG in the dark is completely blocked in *cal1-1* mutant plants, while neither the total starch accumulation nor the dark-induced starch degradation seems to be affected. In CAL1ox plants, MLG is constitutively degraded preventing its accumulation during the day and the content remains very low in all conditions tested. However, no differences were found compared to the control (CAL1ox-negative plants) in the amount of total starch in any of the conditions analyzed here. These observations suggest that CAL1-dependent MLG turnover does not impact starch metabolism under normal conditions.

Taken together, we propose a model where maize plants regulate the accumulation of MLG during the day time by blocking the expression of the CAL1 licheninase. During the night, CAL1 expression is induced/de-repressed to degrade MLG. The lack of obvious developmental or reproductive effects in plants over- or under-accumulating MLG (i.e. *cal1* and *CAL1ox* plants, respectively) grown under the conditions used here does not give further insights into the physiological role of MLG turnover. However, the extended conservation of CAL1-like proteins among diverse grass species suggest an evolutionary advantage of the mechanism of MLG degradation. Indeed, the existence of such precise regulation of MLG content suggests these enzymes have an important biological role.

## DISCUSSION

### MLG degradation

Grass cell walls accumulate large amounts of MLG in different tissues. Synthesis of this polysaccharide is performed by members of the CSLH and CSLF protein families. However, accumulation of MLG is not only affected by the efficiency of its biosynthesis. For example, no direct relationship was found between either the expression of MLG synthase genes or the synthase activity with the level of MLG in barley varieties (Tsuchiya et al., 2005; Burton et al., 2008). Instead, degradation of MLG through specific licheninases is also genetically determined, and has a strong impact on the accumulation of MLG in a particular tissue, developmental stage and growing conditions (Roulin et al., 2002). QTL studies have identified licheninases as important factors to explain the variation in the grain MLG content among diverse barley varieties (Han et al., 1995; García-Gimenez et al., 2019). Licheninase gene expression, protein accumulation, and activity has been found in germinating barley and wheat seeds (Fincher et al., 1986; Litts et al., 1990; Wolf et al., 1991; Lai et al., 1993), suggesting a common mechanism among grasses. Here, we have identified *cal1*, a maize mutant impaired in MLG degradation. *In vitro*, CAL1 exhibits specific licheninase activity, and its overexpression in maize results in a virtual lack of MLG accumulation. Interestingly, the maize genome encodes multiple proteins with predicted licheninase activity belonging to the GH17 and GH1 families, showing high sequence similarity to CAL1. However, our data strongly suggest limited functional redundancy for MLG degradation in maize seedlings, as the MLG content found in *cal1* mutant seedlings is similar to the maximum accumulation measured in the wild-type. Additionally, the four recombinant proteins showing the highest protein sequence similarity to CAL1 failed to degrade MLG in our in vitro activity assays despite the predicted licheninase activity. Although we cannot rule out the possibility that these or other proteins are involved in MLG degradation under specific conditions, developmental stages and/or specific cell-types, our data suggest that CAL1 is not only necessary but sufficient for MLG degradation.

### Regulation of MLG accumulation

MLG accumulation has long been known to be stage-specific, its maximum abundance occurring during phases of rapid elongation and diminished in adult tissues (Carpita, 1996; Kim et al., 2000). In addition, our results show that MLG content also changes during day/night cycles in maize seedlings. Under the conditions tested, maximum MLG abundance is observed at the end of the day and rapidly decreases during the night. This MLG dark-induced degradation of MLG is dependent on *CAL1* expression as it is absent in the *cal1* mutant. Proteins with high sequence similarity to CAL1 can be found encoded in the genomes of many grass species, both cereal and non-cereal, suggesting a conserved mechanism. In fact, many previous observations are in line with the existence of a dark-induced MLG degradation in barley, rice and wheat. For example, CAL1-like protein levels and licheninase activity significantly increase in barley and wheat coleoptiles after prolonged dark incubation periods (Roulin et al., 2001; Roulin et al., 2002). Similarly, MLG content is lower in rice dark-grown coleoptiles compared to those grown in light, coincident with increased licheninase activity (Chen et al., 1999). Our results expand on the idea that MLG may serve as an energy storage polymer, where in young growing tissue MLG accumulates during the day, potentially as a way to store excess carbon derived from photosynthesis, and degraded during the night, providing glucose molecules under conditions of sugar depletion. However, blocking MLG accumulation and turnover as in the *cal1* mutant does not seem to affect the developmental phenotype of the maize seedlings at least under the growth conditions used here. It is likely that based on the amount of MLG that is turned-over per day (around 25 μg/mg AIR) starch seems to be the dominant carbon storage polymer with approximately 100 μg/mg dry weight turnover per day (**Fig. S8**), thus allowing the plant to maintain its growth rate. However, MLG turnover may make a difference in specific circumstances such as plants grown under biotic/abiotic stresses. MLG has also a structural role in some specific cell types such as the vascular tissue (Trethewey et al., 2005; Fincher, 2009; Vega-Sanchez et al., 2015), where it could be protected from licheninase degradation.

### Engineering of MLG content in crops

Composed of (1,3)-β-linked cellotriosyl and tetraosyl units, MLG is efficiently hydrolyzed by glucanases present in commercial enzymatic cocktails (e.g. Accelerase 1500). Thus, the enhanced saccharification yield observed when processing *cal1* plant material is not surprising. However, the increase in MLG content alone (around 20 μg/mg AIR; **Fig. 2**) does not account for the enhanced saccharification yield (around 80 μg/mg AIR; **Fig. 1**). As no other polymer seems to be affected, one possible explanation could be that the high MLG content in *cal1* affects the accessibility of the saccharification enzymes to cellulose. The amount of Glc released in *cal1* walls after mild acid treatment (TFA) is also higher than one would expect based only on the difference in MLG content (33 μg/mg AIR in wildtype plants and 115 μg/mg AIR in *cal1*; **Fig. 1**). It has been hypothesized that MLG might form a gel-like matrix interacting with cellulose and assisting in the proper bundle organization of microfibrils (Fincher, 2009; Smith-Moritz et al., 2015; Kim et al., 2018). As TFA can hydrolyze amorphous cellulose but not crystalline cellulose, one possible explanation could be that the high MLG content in *cal1* somehow alters cellulose-MLG interaction improving the accessibility of the saccharification enzymes and/or altering the crystallinity of cellulose. Also unexpected was the synergistic effect observed when stacking *cal1* with individual mutations affecting lignin (i.e. *bm1* or *bm3*). Lignin negatively impacts lignocellulosic enzymatic hydrolysis in two ways. First, aromatic compounds bind enzymes, removing them from their substrate. Second, self-aggregation of lignin polymers results in a highly hydrophobic composite, reducing the accessibility of the enzymes and the required water to the polysaccharide substrate. Changes in the lignin content or the modification of the monolignol composition alters also the interaction between lignin and other wall polymers (e.g. cellulose). The combination of high MLG content with altered lignin in the *cal1 bm* double mutants results in higher digestibility than the addition of both individual effects. This synergy may be explained by the generation of an overall cell wall architecture that results in better enzyme substrate accessibility, as no additional compositional differences were observed. This combination thus has the potential to improve plant feedstocks for the conversion of lignocellulosic biomass into renewable biofuels and other commodity chemicals.

High digestibility in maize is a desirable trait for the feed industry. In fact, commercial maize *bm3* hybrid lines showing reduced lignin content and increased digestibility are currently being marketed for silage used in dairy production, as these traits have been associated with increased nutrient intake and higher milk production (Oba and Allen, 1999; Coons et al., 2019). High MLG content also enhances nutrient intake in cattle as rumen microbes can hydrolyze MLG (Grove et al., 2006). The *cal1 bm* lines thus have the potential of combining both advantages. Furthermore, MLG content is also a potential target to improve the characteristics of grass crops as feedstocks for food, as MLG is an important source of fiber intake in the human diet. Its uptake has been associated with benefits to gut microbiota and human health, including hypercholesterolemia, diabetes, obesity and cancer (reviewed in Jayachandran et al, 2018).

Multiple approaches aiming at increasing MLG levels in several grass species by over-expressing synthase genes turned out to show only limited success due to associated developmental defects. For example, although CslF6 overexpression results in an increased accumulation of MLG in barley or *Brachypodium*, severe growth problems leading to lethality become apparent. Detailed analyses of these plants showed abnormal vascular development, affecting transport of water and nutrients (Burton et al., 2011; Kim et al., 2015; Fan et al, 2018; Kim et al., 2019).

Our results indicate that maize plants with high MLG content can be obtained by blocking MLG degradation mediated by CAL1. CAL1-deficient plants do not show any obvious developmental or reproductive phenotypes in either greenhouse or field conditions. This approach likely avoids the problems of ectopic deposition of MLG or absence of auxiliary enzymes required for the correct deposition of MLG in the wall as a suggested cause of the ‘vascular suffocation’ in overexpressing MLG-synthase strategies (Burton et al., 2011; Kim et al., 2015). Accordingly, the high MLG content associated with *cal1* does not alter biomass or grain yields. In addition, the disease susceptibility against *U. maydis* in *cal1* seems to be unaffected suggesting that the increase in MLG content may not change the overall cell wall architecture in a way that favors the penetration of specialized wall-degrading fungi. Transference of these traits to other grasses seems possible, as CAL1 is highly conserved in cereals including rice, wheat, barley and millet, and also energy crops as such as miscanthus, switchgrass and sugarcane.

## MATERIAL AND METHODS

### Mapping, plant materials and growth

The *cal1-1* mutant was identified from a F2 mutagenized population generated by ethylmethane sulfonate (EMS) mutagenesis in the A619 inbred (Neuffer 1982; Lewis et al 2014). The F1 plants were self-pollinated and 12 seeds per F2 family were planted in the greenhouse. F2 plants were grown under 16h day/ 8h night photoperiod, temperatures of low 20°C/ high 25.6°C and watered two times per day with 100 ppm M, W, F w 20/20/20 fertilizer (Peters Professional). After two weeks, plants were transferred to a dark room for 20 hours and the second leaf blade collected for cell wall analyses. Only normal-appearing seedlings were analyzed. Putative mutants were transplanted and grown to maturity to cross with a different inbred for mapping purposes *. cal1* was crossed to B73 and then self-pollinated to generate an F2 mapping population. The map position was determined by pooling 30 *cal1*-*1* individuals and a similar number of normal siblings and performing bulked segregant DNA mapping (Michelmore et al., 1991) which identified a locus on chromosome 6. The *cal1* causal locus was narrowed to a region between 141.46MB and 142.89MB of Chromosome 6 (**Fig. S6**). Sequencing of the endoglucanase candidate gene (*GRMZM2G137535*) found within this interval, showed a base pair change consistent with EMS mutagenesis (Primer sequences can be found in **Table S5**). A list of all predicted genes contained in the mapping interval is provided in **Table S4**.

*cal1-2* (identification number PV03 43 H-03) was obtained from the Trait Utility System for Corn (TUSC) collection developed by Pioneer Hi-Bred International (Meeley and Briggs 1995). Primers listed in Supplementary Table 4 were used to genotype these plants.

*bm1* and *bm3* mutants were kindly provided by the Maize Genetics Coop (Catalog#515D and 408E, respectively).

A *cal1-1* dCAP marker was used to identify the mutation in segregating populations. cal1MmeIF and R primers were used to amplify a 862bp amplicon. PCR fragments were treated with *MmeI* restriction enzyme (Thermo) according to manufacturer’s instructions. Wildtype-derived PCR amplicons resulted in three bands of 330, 316 and 127bp, while *cal1-1* mutant samples showed only two bands (647 and 127bp) due to a G-A nucleotide change in one of the *MmeI* restriction sites.

### Generation of CAL1 overexpression lines

For the generation of *CAL1* overexpression lines, full-length coding sequences from *CAL1* was obtained from GeneArt (www.thermofisher.com) and then subcloned into the *pUbi-BASK* vector using *BamHI* and *SpeI* restriction enzymes. The expression cassette including *UBI5* promoter, *UBI5’* Intron, *CAL1* coding region and a *NOS* terminator was excised using *EcoRI* and *HindIII* restriction enzymes and cloned into binary *pTF101* vector (Paz et al., 2004; Addgene plasmid #134770; http://n2t.net/addgene:134770; RRID: Addgene_134770).

Hi Type II hybrid immature zygotic embryos were transformed with the *pTF101:CAL1* construct via Agrobacterium-mediated transformation along with at the Iowa State University Plant Transformation Facility (Frame et al., 2002 and http://agron-www.agron.iastate.edu/ptf/service/agromaize.aspx). T0 transformant plantlets were assayed for transgene presence by screening for BAR marker gene-conferred resistance. A 5cm diameter circle was drawn on an adult leaf (usually leaf 10) with a marker and applying 20ul/mL Finale® (11% ammonium gluphosinate) with 1μl/ml Tween20^®^ detergent with a cotton swab. Expression of resistance or susceptibility was studied by visual inspection after 5-7 days. Plants were genotyped for the presence of the transgene using primers newPTF101R and CL 391. To verify transgene expression, RNA was extracted from second leaves of 3 plants showing a positive genotyping result for each independent line and reverse-transcribed. Two pools of 3 plants negative for the transgene were used as controls. Quantitative RT-PCR was used to test relative expression in the different lines by amplifying the resulting cDNA using primers CL 407/CL 408.

### Cell wall analyses

Plant tissues were lyophilized in a ScanVac CoolSafe Freeze-dryer (Labogene) for 48h and homogenized in 2mL screw cap tubes containing two 5mm steel beads for 2 min at 30 Hz in a MM400 mixer mill (Retsch Technology). Preparation of dAIR and determinations of crystalline cellulose content, total lignin content, and monolignol composition were performed as indicated in Foster et al., 2010a, 2010b. For the biomass monosaccharide composition to screen for maize mutants, a Shimadzu Prominence HPLC system equipped with a refractive index detector was used to measure glucose, xylose, and arabinose in the TFA hydrolysates. Samples were separated with a Phenomenex RezexTM RFQ-Fast Acid H^+^ (8%) ion exchange column (100 × 7.8 mm) and a Bio-Rad Cation H guard column (30 × 4.6 mm) with 5 mM sulfuric acid as the mobile phase and a flow rate of 1.0 mL min^−1^ at 55°C for 5 min. The determination of total wall acetate content was performed as described in Ramirez et al., 2018.

For saccharification assays, 1mg dAIR was incubated in the presence of 0.5μL of Accelerase 1500 (Genencor) in 0.1M citrate buffer pH4.5 and 0.15mM NaN_3_ in a 1.2mL final volume reaction (Santoro et al., 2010). One 5mm steel bead was added and reactions were incubated at 50°C for 20h shaking at 250rpm. Released Glc was measured in a YSI 2900 biochemistry analyzer (Xylem Inc.) following manufacturer’s instructions.

Determination of the MLG content was performed using a mixed-linkage beta-glucan assay kit (K-GLU kit, Megazyme) with modifications. In brief, two milligrams of AIR were resuspended in 200 μL 20mM NaPO_4_ buffer pH6.5 and incubated 5 minutes at 100°C. After cooling down to room temperature, 20 μL Licheninase [specific, endo-(1-3); (1-4)-β-D-glucan 4-glucanohydrolase; Megazyme] were added and the mix incubated at 40°C for 1h. After centrifugation 10min at 12000 rpm, 10 μL of the supernatant containing MLG-derived oligosaccharides was mixed with 10 μL β-Glucosidase (Megazyme) and incubated 15min at 50°C. After adding 300 μL GOPOD reagent, the mix was incubated 20 min at 50°C. OD_510_ was measured and the Glc content in the samples was calculated using a Glc standard curve. Glc content obtained from samples not treated with licheninase but treated with β-glucosidase was subtracted.

For the determination of MLG fine structure, 2mg of AIR were treated with 4M KOH for 4 h under constant shaking. Samples were centrifuged for 10 min at 12000 rpm and supernatants transferred to a new tube and pellets washed three times with 1.5 mL water. Supernatants from all steps were combined resulting in the 4 M KOH-soluble fraction and neutralized with concentrated HCl. KOH-soluble fraction was then digested with licheninase (Megazyme) as described before. The supernatant obtained was then filtered through 0.45 μm PTFE filter units (Millipore), diluted 1:10, and injected in an ICS-3000 Dionex chromatography system (Dionex, Sunnyvale, CA, USA) equipped with a CarboPac PA200 anion-exchange column. Identification and quantification of the present cello-oligosaccharides (DP3-DP6) was performed using cellotriose, cellotetraose, cellopentaose and cellohexaose commercial standards (Megazyme).

For the Saeman hydrolysis of licheninase-resistant residues, samples were dried in a Speed Vac before adding 175μL of 72% (w/w) H_2_SO_4_. Samples were incubated at room temperature for 45 minutes. Glc content of the supernatant was determined by anthrone assay (Dische, 1962). Briefly, 10μL sample was mixed with 90 μL water and 250μL anthrone reagent [0.2% (w/w) anthrone (Sigma) in concentrated H_2_SO_4_]. Glc concentration was quantified by reading the absorbance at 620nm.

### Starch analysis

Starch content was determined using the Total Starch Assay Kit (Megazyme) with the following modifications. 1mg AIR was resuspended in 100μL thermostable amylase and incubated 15 min at 100°C. Samples were allowed to cool down and 10μL supernatant was mixed with 10μL α-glucosidase and incubated 30 min at 50°C. Glc content was determined using the GOPOD method as described above.

### Production and purification of His-tagged proteins

Full-length coding sequences from CAL1, *cal1* and maize paralogs were obtained from GeneArt (www.thermofisher.com). A 3’ Histidine tag coding sequence was added using primers listed in **Table S5** and the resulting PCR products cloned in TOPO/TA vector according to manufacturer’s instructions (www.thermofischer.com). His tag-containing fragments were excised from pTOPO/AT and cloned into pBART27 (Stintzi and Browse, 2000) vector using XhoI and BamHI restriction sites.

pBART27 constructs were transformed into *Agrobacterium tumefaciens* strain GV3101. Transient transformation of *N. benthamiana* was performed as described in Ramirez et al., 2013. Infiltrated leaves were collected after 3 days and immediately frozen in liquid nitrogen. Frozen leaves were milled to a fine powder using mortar and pestle. Extraction buffer [50mM sodium phosphate buffer, 1M NaCl, pH8, 10 μL/mL of Protease Inhibitor Cocktail (Sigma) and 0.14 μL/mL of β-mercaptoethanol] was added in a 1:3 ratio (powder/buffer) and incubated 1h at 4°C in a rotatory wheel. After filtering through 0.45μm filter units (Merck), His-tagged proteins were washed and purified using the Amicon Pro System (Merck) according to manufacturer’s instructions.

### Activity assays

Glycanase activity assays were performed in 20mM sodium citrate buffer pH4.5 at 37°C for 1h using 2μL purified recombinant proteins and 25μg substrate. 150 μL water were added and reactions heat-inactivated at 100°C for 10 min. The amount of reducing ends generated was determined using the PAHBAH assay (Lever et al., 1973). Briefly, PAHBAH reagent was prepared by dissolving 5% (w/v) 4-hydroxybenzoic acid hydrazide in 0.5N HCl and mixed 1:5 (v/v) with 5M NaOH. 750 μL PAHBAH reagent were added to the reaction mix and incubated 5 min at 100°C. 200 μL aliquots were then transferred to 96-well UV-plates and Abs_410_ determined.

All substrates, (1,3-1,4)-β-D-Glucan from barley flour, laminarin and were obtained from Megazyme.

### Gene expression analyses

Total RNA was extracted from pools of 3-5 second leaves using the Plant RNAeasy kit (Qiagen) using manufacturer’s instructions. DNAse-treated RNA (Turbo DNAse, Ambion) was used for cDNA synthesis (iScript, Biorad). Semi-quantitative RT-PCR was performed on 0.8μL of cDNA resulting from reverse transcription of 500ng of RNA. Specific primers were used to amplify CAL1 and GAPDH as housekeeping control using Phusion Taq polymerase (Thermo Scientific). Reaction conditions and annealing temperatures followed the manufacturer’s suggestions. Products were resolved using 1% agarose gels and visualized using 0.5μg/mL ethidium bromide. RT-qPCRs were performed in a Biorad CFX96 system using recommended parameters and data analyzed by the CFX Master Software. Actin2 or GAPDH were used as housekeeping genes. Primer sequences can be found in **Table S5**.

### Sequence alignments and Phylogeny

Protein sequences were obtained from the Maize Genetics and Genomics Database (www.maizedgb.org). Protein alignments were performed using Clustal Omega software with default settings (Sievers and Higgins, 2018). Phylogenetic trees were constructed by using the Phylogeny.fr web service (Dereeper et al., 2008).

### *U. maydis* infection assay

The sensitivity of the *cal1-1* line to smut fungal infection was tested in established seedling infection assays as described before (Bösch 2016). The solopathogenic *U. maydis* strain SG200 (Kämper 2006) was grown to an OD600 of 0.8 in complete medium, washed three times with H2O, and resuspended to an OD600 of 3 in H_2_O. The cell suspension was injected into 7-day-old maize seedlings. Plants at 7 dpi and 14 dpi were scored for symptom formation. Categories for disease rating were the following: 1, no symptoms; 2, chlorosis; 3, anthocyanin accumulation; 4, small tumors (<1 mm); 5, medium tumors (1-5 mm); and 6, heavy tumors (>5 mm) associated with bending of the stem.

### Yield determinations

Seeds of *cal1/cal1* mutants in an inbred A619 background were crossed to B73 and Mo17 to obtain wild-type (*Cal1/cal1*) B73/A619 and Mo17/A619 hybrids, respectively. All three entries were planted in Urbana in summer 2011 in 4m rows spaced 0.76 m apart with 112 kg N/ha applied prior to planting. Each of the three entries was planted in 8 replicates, for a total of 24 rows. Seeds were hand-planted every 15 cm. At grain maturity, a single representative plant from each row was harvested, wet grain and wet biomass were weighed separately, and a weighed subsample of grain and biomass was oven-dried at 80C for 3 days and re-weighed to determine moisture content.

## Acknowledgements

We would like to thank Grace Kayser, Jiayin Shum, Yadanar Htike and Cesar Morphin for assistance in mapping *cal1*, Nathalie Bolduc, George Chuck and Devi Santosh for primer design, Vincent Wu for the analysis of CAL1ox saccharification, Anand Narayanan for help characterizing the *cal1-2* allele and Katharina Lufen for technical support on the cell wall analyses.

**List of author contributions:** SH, MP and VR conceived and planned the experiments. FJK and CL identified, characterized and mapped *cal1*. MK and BMK characterized protein activities, CAL1ox lines and *cal1bm* double mutants. CR carried out the seedling growth measurements. PJB analyzed plant yield in a field trial. PH and VG performed the *U. maydis* infection experiments. VR and MP wrote the manuscript with input from all authors.

**Funding information:** This work has been funded by Germany’s Federal ministry of education and research (BMBF) grant “Cornwall”, 031B0193A to MP. Additional funding was provided by the Deutsche Forschungsgemeinschaft (DFG, German Research Foundation) under Germany’s Excellence Strategy – EXC 2048/1 – Project ID: 390686111 to M.P., Marie Curie PIOF-GA-2013-623553 to V.R., and USDA-ARS CRIS 2030-21000-051-00D to SH.

## Supplementary Figure legends

**Fig S1.** *cal1* plants exhibit normal growth habit.

**Fig S2.** cal1 plants exhibit normal growth habit in the field.

**Fig S3.***U. maydis* infection of *cal1*

**Fig S4.** Fine-mapping of *cal1* mutation on Chromosome 6.

**Fig S5.** A maximum likelihood protein sequence phylogenetic tree of CAL1

**Fig S6.** Overexpression of CAL1 in maize.

**Fig S7.** Phylogenetic tree of maize proteins with similarity to CAL1

**Fig S8.** Dark-induced MLG and starch degradation

**Table S1.** Cell wall composition of various tissues of the maize *cal1* mutant.

**Table S2.** Total weight and moisture content in *cal1* maize mutant plants.

**Table S3.** Lignin content and composition in *cal1 bm* double mutants.

**Table S4.** Primers used in this study

**Table S5.** List of genes included in the *cal1* mapping interval.

